# Controlled In Vitro Characterization of the Dynamic Response of Continuous Glucose Monitoring Systems: Adaptation of a Programmable Flow Platform and Decomposition of Dynamic Error

**DOI:** 10.64898/2026.08.12.743851

**Authors:** Elena V. Khoroshun, V. Kozlov, Igor V. Ivanov, Kuvat T. Momynaliev

## Abstract

**Background:** Continuous glucose monitoring (CGM) systems are used not only for retrospective assessment of the glycemic profile but also for real-time decision-making, including automated insulin delivery. Accordingly, CGM performance characterization must capture not only the agreement of individual paired values but also the system’s ability to reproduce the direction, rate, amplitude, and shape of glucose concentration change. Summary metrics, most notably MARD, cannot establish whether an observed deviation reflects an error in the formation of the test profile itself, a constant sensor offset, amplitude compression, a change in response rate, temporal misalignment, or hysteresis.

**Objective:** To adapt a programmable flow-based in vitro platform for the separate assessment of the experimentally delivered glucose profile and the dynamic response of CGM systems, and to propose a set of metrics that decomposes dynamic error into its components.

**Methods:** GLU profiles were generated by programmable mixing of solutions at a constant total flow rate of 2 mL/min. Actual GLU concentration was independently measured with a SUPER GL2 glucose analyzer. Four static levels, three repeats of a 5.5→12.0→5.5 mmol/L profile, three repeats of a 6.0→3.0→6.0 mmol/L hypoglycemic profile, three 5.0→15.0→5.0 mmol/L profiles at different rates, one complex 4→18→3→12→5.5 mmol/L profile, and two proof-of-concept sensor experiments at 100- and 200-min transitions were investigated. Dynamic response was characterized by bias, MAE, RMSE, MARD, amplitude transfer coefficient K_A, rate transfer coefficients K_up and K_down, normalized shape RMSE, residual shift, and hysteresis loop area.

**Results:** At the static levels, measured GLU exceeded the programmed value by 0.234–0.780 mmol/L. In the repeated 5.5→12.0→5.5 profiles, the ratio of actual to programmed rate was 0.978–1.083 on the rising phase and 0.987–1.157 on the falling phase, while the amplitude transfer coefficient was 0.967–1.066. In the hypoglycemic profile, minimum GLU was 2.55– 2.96 mmol/L, and time below 3.0 mmol/L was 15.2–72.6 min. The measured rates of 0.0519, 0.1045, and 0.2027 mmol/L/min preserved the intended ratio of approximately 1:2:4. In the complex profile, the programmed plateau of 18 mmol/L was not reached: mean measured GLU was 16.20 mmol/L. For CGM-A, K_A was 0.682 and 0.650, and K_up/K_down were 0.666/0.730 and 0.634/0.626; the corresponding values for CGM-B were 1.228 and 1.128, and 1.564/1.328 and 1.276/1.145. Hysteresis loop area differed 5- to 10-fold between the two sensor responses, exceeding an order of magnitude at the 100-min transition.

**Conclusion:** The programmed concentration should be treated as a control setpoint, rather than as a reference measurement. The “programmed trajectory — measured glucose — CGM output” cascade first allows quantitative assessment of the agreement between the programmed and actually realized profile and only then separate characterization of sensor response.

Decomposition of dynamic error into amplitude, rate, shape, and hysteresis components reveals differences that a single MARD value or correlation coefficient cannot capture.

## Introduction

Continuous glucose monitoring (CGM) systems have become a central element of modern diabetes care. They provide a continuous time series from which current glucose level, direction and rate of change, time in range, variability, and hypo-/hyperglycemia risk are derived (Danne et al., 2017; Battelino et al., 2019). When CGM is used in automated insulin delivery systems, what matters is not only static accuracy but also the sensor’s ability to reproduce a rapidly changing profile promptly and proportionally. Error at steady-state concentration and error during rapid change can have different mechanisms and different clinical significance (Kovatchev et al., 2004; Klonoff et al., 2024).

A CGM reading is the product of a multistage process. Under clinical conditions it is shaped by glucose transport between blood and interstitial fluid, local perfusion, membrane and enzyme-layer properties, the electrochemical response, temperature sensitivity, calibration, algorithmic filtering, update frequency, and display rules (Rebrin and Steil, 2000; Cengiz and Tamborlane, 2009; Schmelzeisen-Redeker et al., 2015). The observed discrepancy between CGM and the comparator method therefore results from the superposition of physiological, analytical, and algorithmic components. Clinical studies are necessary to assess overall safety and effectiveness, but they do not always allow the technical source of dynamic error to be identified.

The most widely used summary accuracy metric for CGM remains the mean absolute relative difference (MARD). Its simplicity has driven wide adoption, but MARD depends on the concentration distribution, the number and temporal density of pairs, the accuracy of the comparator method, synchronization rules, and the proportion of observations across different GLU ranges (Kirchsteiger et al., 2015; Pleus et al., 2017; Heinemann et al., 2020; Vigersky and Shin, 2024). The same MARD value can arise from fundamentally different error types: a constant negative bias, proportional overestimation, amplitude compression, delay, or phase-dependent discrepancy. High correlation is likewise not proof of agreement, since a sensor can accurately reproduce the shape of a profile while exhibiting a pronounced systematic or proportional error (Clarke and Kovatchev, 2007; Freckmann et al., 2023).

Current approaches to CGM evaluation are gradually shifting from a single integral number toward a multicomponent characterization. Continuous glucose-error grid analysis links analytical error to clinical risk (Kovatchev et al., 2004), while CG-DIVA describes the distribution of deviations and accuracy variability within and between sensors (Eichenlaub et al., 2024). These methods extend the interpretation of point accuracy but do not by themselves explain how a sensor transmits amplitude, rate, and direction of change over time.

The dependence of CGM accuracy on the rate of glucose change has been demonstrated in clinical studies using induced glycemic swings: MARD can increase during rapid changes, and the degree of deterioration differs between systems (Pleus et al., 2015). Controlled in vitro experiments make it possible to exclude physiological inter-compartmental transport and thereby isolate the intrinsic sensor response, along with the influence of filtering and update frequency. Davey et al. (2010) showed that the system’s own lag can contribute markedly to the discrepancy at variable rates. Similarly, experimental work demonstrates that algorithmic rate-of-change limiting can create an apparent lag even in the absence of a physiological gradient.

However, the laboratory test bench is itself a measurement system and may produce a profile that differs from the intended one. Errors in flow dosing, incomplete mixing, dead volume, thermal conditions, adsorption, transport delay, and tubing properties can all alter the actual amplitude and shape of the delivered signal. Consequently, comparing CGM output directly against the programmed trajectory risks misattributing to the sensor a deviation that in fact arose at the platform level. The quality of the comparator method and temporal synchronization critically affect the resulting accuracy metrics (Kirchsteiger et al., 2015; Freckmann et al., 2023).

Pfützner et al. (2024) demonstrated the feasibility of controlled laboratory dynamic testing of CGM systems. Further development of this approach calls for an analytical hierarchy in which the programmed trajectory serves as the control setpoint, the delivered glucose profile is independently verified by a comparator method, and the sensor is evaluated only against the measured GLU. Such a hierarchy allows the reproducibility of the platform and the dynamic properties of the sensor to be assessed separately.

The clinical relevance of CGM dynamic accuracy is growing as automated insulin delivery (AID) systems become more widespread, since in these systems the insulin dosing algorithm relies directly on the current sensor reading and trend without an intermediate user check (Boughton and Hovorka, 2024). In such systems, an error that arises specifically during rapid GLU change, rather than at steady state, can translate directly into an incorrect dosing decision. This is also reflected in the evolution of regulatory accuracy-assessment tools: the revised Diabetes Technology Society Error Grid has been supplemented with a separate Trend Accuracy Matrix, which for the first time formalizes the clinical risk of errors specifically in the assessment of glucose rate and direction of change, rather than the point value alone (Klonoff et al., 2024). The emergence of such tools underscores the need for laboratory methods capable of quantitatively and reproducibly characterizing sensor dynamic response already at the bench-testing stage, prior to clinical evaluation.

In the present study we adapted a programmable flow platform and generated a set of static, repeated, hypoglycemic, rate-controlled, and complex profiles. We hypothesized that sequential assessment along the “programmed trajectory — measured glucose — CGM output” cascade would allow the observed error to be decomposed into a profile-generation component and a sensor-response component. The aim of the study was to verify the platform’s capabilities and to demonstrate a set of metrics characterizing systematic bias, amplitude and rate transfer, normalized shape error, residual shift, and hysteresis.

## MATERIALS AND METHODS

### Design and Analytical Hierarchy

The study comprised two levels of analysis. At the first level, the programmed concentration was compared against the GLU concentration actually measured by the SUPER GL2 analyzer. At the second level, CGM readings were compared exclusively against the time series of measured GLU. This approach follows the principle that the quality of the comparator method and temporal synchronization substantially affect CGM evaluation (Kirchsteiger et al., 2015; Freckmann et al., 2023).

### Platform and Reference Analyzer

GLU profiles were generated by programmable mixing of two solutions of identical matrix: solution A with a nominal glucose concentration of 20.0 mmol/L, and solution B with no added glucose. The working medium was phosphate-buffered saline (PBS) at pH 7.2±0.2. Total flow rate was held constant at 2.0 mL/min, while the time-varying split between streams A and B produced the required static levels, linear transitions, and multiphase profiles. The calculated concentration derived from the flow ratio was treated as the control setpoint; actual GLU was confirmed by independent measurement.

Prior to the sensor experiments, platform qualification included verification of spatial homogeneity within the macrofluidic manifold. This qualification was performed as a separate protocol using its own set of programmed levels (3.5, 5.5, 11.0, and 20.0 mmol/L), which did not coincide with the levels used in the static bias experiment reported below. At each level, GLU was measured in randomized order across all six predefined control positions (S1–S6), with four replicates per position (n = 24 per level). Randomization of the sampling order minimized systematic bias associated with sampling sequence and allowed discrimination between true spatial concentration gradients and ordinary experimental variability. One-way ANOVA and the Kruskal–Wallis test showed no significant effect of position at any level (3.5 mmol/L: F = 1.29, p = 0.313, Kruskal–Wallis H = 6.64, p = 0.249; 5.5 mmol/L: F = 0.62, p = 0.686, H = 3.38, p = 0.642; 11.0 mmol/L: F = 0.52, p = 0.758, H = 2.41, p = 0.790; 20.0 mmol/L: F = 1.34, p = 0.294, H = 7.23, p = 0.204); the range of position means was 0.09–0.80 mmol/L across the four levels. The manifold was therefore considered spatially homogeneous under the experimental conditions of the present study.

Experiments were performed on a bench built for laboratory in vitro testing of CGM sensors, comprising a programmable Agilent 1260 Infinity III Flexible Pump quaternary pump, two reservoirs of working solution of identical matrix, a degassing system, supply and return lines, a macrofluidic flow manifold, a heating system with a remote feedback sensor, a peristaltic pump in the drain line, a control-sample port, a SUPER GL2 glucose analyzer, and a data-acquisition system. The pump was used to independently dose solutions A and B and continuously mix them at a constant total flow rate of 2.0 mL/min. Solution A contained GLU at a nominal concentration of 20.0 mmol/L, while solution B contained no added GLU; both solutions were prepared in the same PBS matrix, which minimized changes in ionic composition and pH as the flow ratio was varied.

The flow unit was a macrofluidic manifold manufactured from polypropylene, weighing approximately 0.4 kg, with overall dimensions of 150×150 mm and an external corner radius of R10. An open S-shaped flow channel of nominal width 2 mm and depth 10 mm was formed in the top surface, as shown in the manifold’s dimensional drawing (Figure 1c). The calculated internal channel volume was determined by the three-dimensional design model and was not separately given on the available two-dimensional drawing. The working medium successively traversed two turns of the channel from the inlet to the outlet port, allowing several sensors to be placed within a single thermostatically controlled zone and their readings to be compared at a common input GLU concentration.

**Figure 1.**
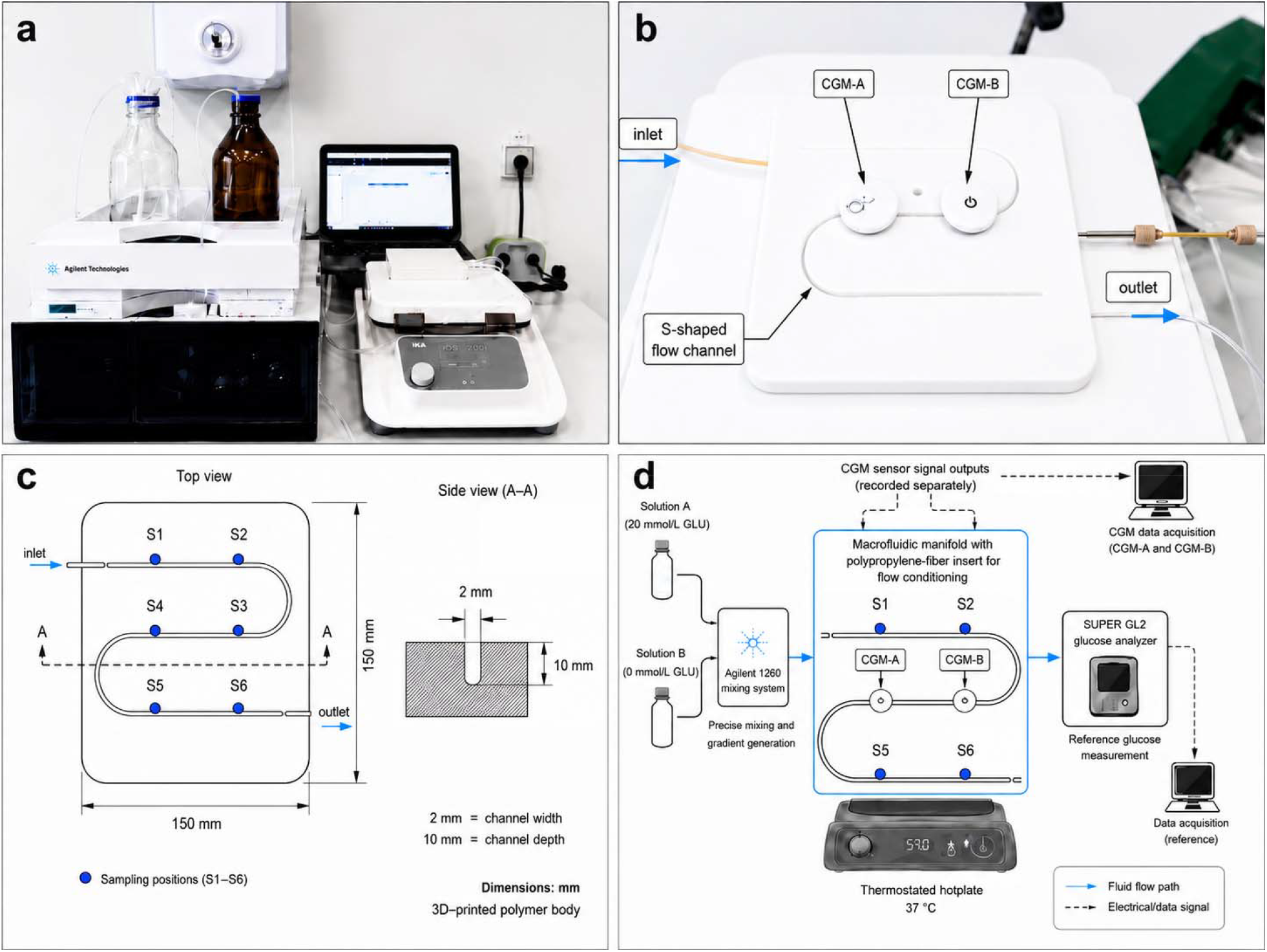
Test setup, macrofluidic manifold design, and analytical cascade. (a) Overall view of the test bench: a programmable Agilent 1260 Infinity III pump, reservoirs of working solutions A and B, and the data-acquisition system. (b) The macrofluidic manifold with CGM-A and CGM-B sensors installed in the working zone; the inlet and outlet ports and the S-shaped flow channel are indicated. (c) Dimensional drawing of the manifold: top view with control positions S1–S6 along the channel, and side view (section A–A) showing channel width (2 mm) and depth (10 mm) within an overall body size of 150×150 mm; the manifold body was manufactured by 3D printing from polypropylene. (d) Schematic of the analytical chain: programmed mixing of solutions A and B, passage of the medium through the manifold with sensors CGM-A and CGM-B, independent measurement of GLU by the SUPER GL2 analyzer, and parallel recording of CGM sensor data and the reference measurement.

A polypropylene fiber insert was placed in the sensor-installation zone. In preliminary technical trials it was used to promote laminar flow; in the present work it was functionally regarded as a passive flow-conditioning element that reduces local jetting, evens out medium distribution around the sensing elements, and lessens the influence of bubbles and local inhomogeneities.

Reynolds number calculation and independent quantitative verification of the laminar regime were not performed within the scope of this study.

To describe the spatial arrangement of the sensors and to assess condition uniformity, six sequential control positions S1–S6 were defined along the S-shaped channel in the direction of flow. S1 and S2 corresponded to the inlet limb of the channel, S3 and S4 to the intermediate section following the first and second changes in flow direction, and S5 and S6 to the outlet limb. These designations were geometric control positions on the manifold rather than separate measurement channels. Spatial homogeneity across these six positions was formally verified by the randomized qualification experiment described above, which also included verification of the temperature distribution within the working region.

The manifold was mounted on a ceramic HP500-Pro Hotplate (DLAB Scientific Co., Ltd., Beijing, China) heating surface. A remote feedback-loop sensor was inserted into the manifold’s central bore and advanced to the lower part of the S-shaped flow channel. Time for the manifold to reach the set temperature of 37 °C was approximately 12 min. The spatial temperature profile was additionally verified with a calibrated Fluke 5411 B digital temperature meter, whose thermocouple was placed directly in the flow of working medium and held in place until readings stabilized. The working region was defined as the middle section of the S-shaped channel, in which the mobile-phase temperature remained within ±2 °C of the set value. Sensors were placed precisely within this thermally stable zone. This temperature regime (37 °C) was maintained without exception throughout all experiments of the present study, including the platform’s static and dynamic profiles, the sensor stage, and the complex multiphase profile. Actual GLU concentration was determined with a SUPER GL2 glucose analyzer (Dr. Müller Gerätebau GmbH, Freital, Germany) at a control point after the working medium had passed through the test unit. According to the manufacturer’s operating manual, the measurement range for molar glucose concentration is 0.5–50 mmol/L, and the coefficient of variation for a series of 20 measurements at a level of 12 mmol/L is below 1.5%. The local distributor additionally registered the instrument in the Russian register of measuring instruments under a simplified type description (Rosstandart, Registration No. 74068-19; verification procedure No. MP 051.D4-18), which states a narrower range (4–30 mmol/L) and a permissible relative error of

±10%; these figures reflect the scope of that particular local registration rather than the instrument’s analytical performance as stated by the manufacturer. The resulting time series was used to verify the actually realized profile and as the comparison series for analyzing CGM readings. Where sampling frequencies differed, measured GLU values were linearly interpolated to the sensor’s timestamps only within the observed range, without extrapolation.

### Experimental Profiles

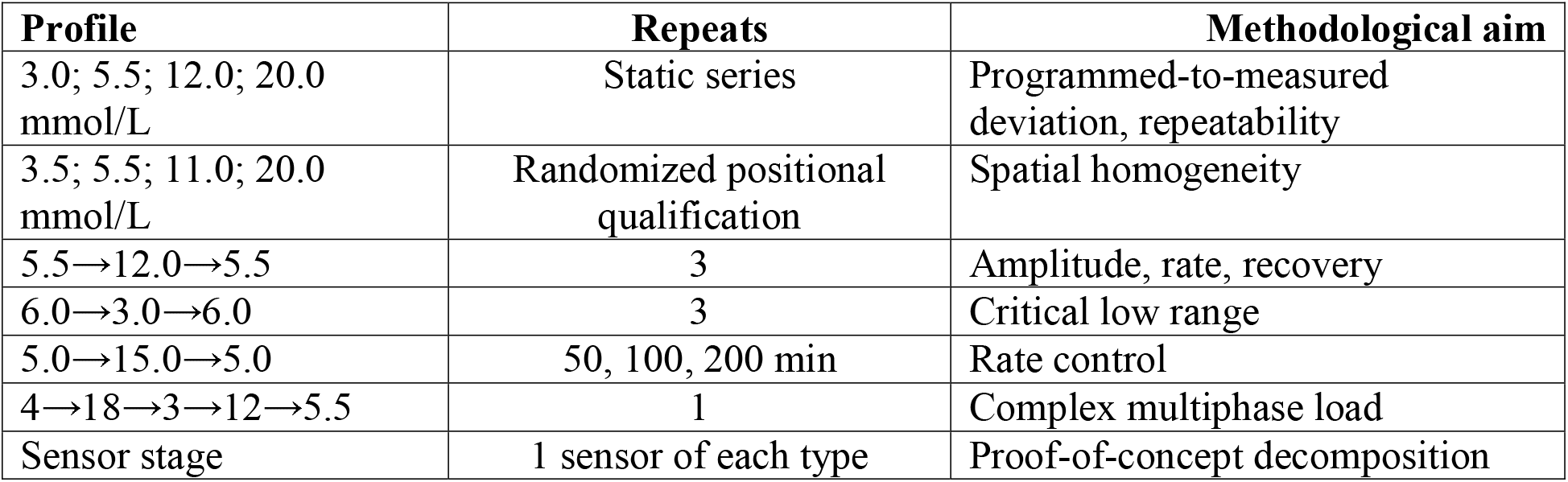

| Profile | Repeats | Methodological aim |
| --- | --- | --- |
| 3.0; 5.5; 12.0; 20.0 mmol/L | Static series | Programmed-to-measured deviation, repeatability |
| 3.5; 5.5; 11.0; 20.0 mmol/L | Randomized positional qualification | Spatial homogeneity |
| 5.5→12.0→5.5 | 3 | Amplitude, rate, recovery |
| 6.0→3.0→6.0 | 3 | Critical low range |
| 5.0→15.0→5.0 | 50, 100, 200 min | Rate control |
| 4→18→3→12→5.5 | 1 | Complex multiphase load |
| Sensor stage | 1 sensor of each type | Proof-of-concept decomposition |

### Data Processing and Metrics

Values of GLU measured by the SUPER GL2 analyzer were interpolated only within the observed time range to the timestamps of each sensor; extrapolation was not applied. Consecutive points of the time series were not treated as independent replicates. The primary platform metrics were programmed-to-measured deviation, the ratio of measured to programmed rate, profile RMSE, and recovery. For the sensor stage, we calculated bias, MAE, RMSE, MARD, the amplitude transfer coefficient K_A, the rising- and falling-rate transfer coefficients K_up and K_down, normalized shape RMSE, residual shift, update frequency, and hysteresis loop area. Correlation was not interpreted as evidence of method agreement (Kovatchev et al., 2004; Freckmann et al., 2023).

### Calculation of CGM Dynamic Response Metrics

For each profile, an initial plateau, a rising phase, an upper plateau, a falling phase, and a final plateau were defined. Phase boundaries were set from the programmed trajectory and applied identically to the measured-GLU series and to the CGM readings.

### Time-series synchronization

Because the SUPER GL2 analyzer and the CGM systems recorded values at different frequencies, measured GLU values were linearly interpolated to the timestamps of each sensor. Interpolation was performed only between adjacent actually measured points; extrapolation beyond the observed range was not applied.

The error of an individual synchronized pair was defined as e_i = CGM_i − GLU_i, where CGM_i is the CGM reading and GLU_i is the GLU value measured by the SUPER GL2 analyzer and interpolated to the same timestamp. Mean systematic bias was calculated as Bias = (1/n)Σ(CGM_i − GLU_i); mean absolute error as MAE = (1/n)Σ|CGM_i − GLU_i|; root-mean-square error as RMSE = √[(1/n)Σ(CGM_i − GLU_i)^2^]. MARD was calculated as (100%/n)Σ|CGM_i − GLU_i|/GLU_i over all synchronized pairs with positive GLU_i. These metrics characterize, respectively, the direction of systematic error, the mean absolute error magnitude, the contribution of large deviations, and the relative error of the CGM system. The amplitude transfer coefficient K_A was calculated as the ratio of the amplitude actually reproduced by the sensor to the amplitude of the GLU change independently measured by the SUPER GL2 analyzer: K_A = ΔCGM/ΔGLU, where ΔCGM = mean(CGM_upper) − mean(CGM_baseline) and ΔGLU = mean(GLU_upper) − mean(GLU_baseline). For the full cycle, the mean values of the initial and upper plateaus were used; for separate analysis of the rising and falling phases, amplitude was determined relative to the corresponding boundary plateaus. A value of K_A = 1 corresponds to proportional amplitude transfer, K_A < 1 to compression, and K_A > 1 to amplification.

The rate transfer coefficients indicate how quickly the CGM sensor reproduces the actual rise or fall of GLU relative to the SUPER GL2 analyzer. Rate was defined as the slope of the linear regression of concentration on time during the corresponding phase. K_up = v_CGM,up/v_GLU,up. For the falling phase, absolute values of the negative slopes were used: K_down = |v_CGM,down|/|v_GLU,down|. A value near 1 corresponds to reproduction of the actual rate; values below 1 point to smoothing or slowing, and values above 1 to amplification of the rate of change.

Normalized RMSE characterized the difference in the shape of the time profiles after removing constant offset and amplitude differences. It was calculated by normalizing each series relative to its baseline level and amplitude: X_norm(t) = [X(t) − mean(X_baseline)]/[mean(X_upper) − mean(X_baseline)]. RMSE was then calculated between the normalized CGM and GLU series. The hysteresis loop area characterized the dependence of CGM readings on the direction of GLU change: whether sensor readings differ at the same GLU value during the rising and falling phases. Hysteresis was assessed in “measured GLU — CGM output” coordinates. The rising and falling branches were analyzed separately; the CGM signal was interpolated onto a common GLU grid spanning the range represented in both the rising and falling data. Area was calculated by the trapezoidal method as A_hyst = ∫|CGM_up(GLU) − CGM_down(GLU)|dGLU, with units of (mmol/L)^2^.

Residual shift characterized the completeness of the sensor’s return to its initial state after completion of a dynamic cycle and was calculated as the difference between mean CGM ‐ GLU error on the final plateau and on the initial plateau. Uniform averaging windows were used for all sensor experiments: the first 20 min of the experiment for the initial plateau, and the last 20 min of the available time series for the final plateau. Median update interval was determined from the differences between consecutive sensor timestamps. Signal loss was defined as the absence of a recorded value for an interval exceeding the system’s expected update interval.

## RESULTS

The experimental program covered several complementary levels of validation. Static series were used to evaluate the correspondence between the programmed and actually measured GLU level. Repeated 5.5→12.0→5.5 mmol/L profiles characterized the reproducibility of shape, rate, and amplitude. The hypoglycemic profile allowed assessment of the depth and duration of time spent in the critically low range. The 5.0→15.0→5.0 mmol/L series with three transition durations tested rate control at constant amplitude. The complex multiphase profile combined several levels and directions of change. The sensor stage was performed as a proof-of-concept: one sensor of each type was investigated at 100- and 200-min transitions; these data were used to demonstrate the analytical decomposition rather than to draw conclusions about the reproducibility of specific commercial models.

### Static Levels and Positional Uniformity

At the programmed levels of 3.0, 5.5, 12.0, and 20.0 mmol/L, mean measured GLU values were 3.282, 5.734, 12.28, and 20.78 mmol/L, respectively. Measured GLU exceeded the programmed setpoint by +0.282, +0.234, +0.280, and +0.780 mmol/L, respectively. Because these experiments were performed during the initial stage of platform qualification, the programmed flow ratios were regarded as nominal setpoints, whereas the experimentally measured GLU concentration was used as the reference concentration for all subsequent analyses of CGM performance (Figure 2a).

**Figure 2.**
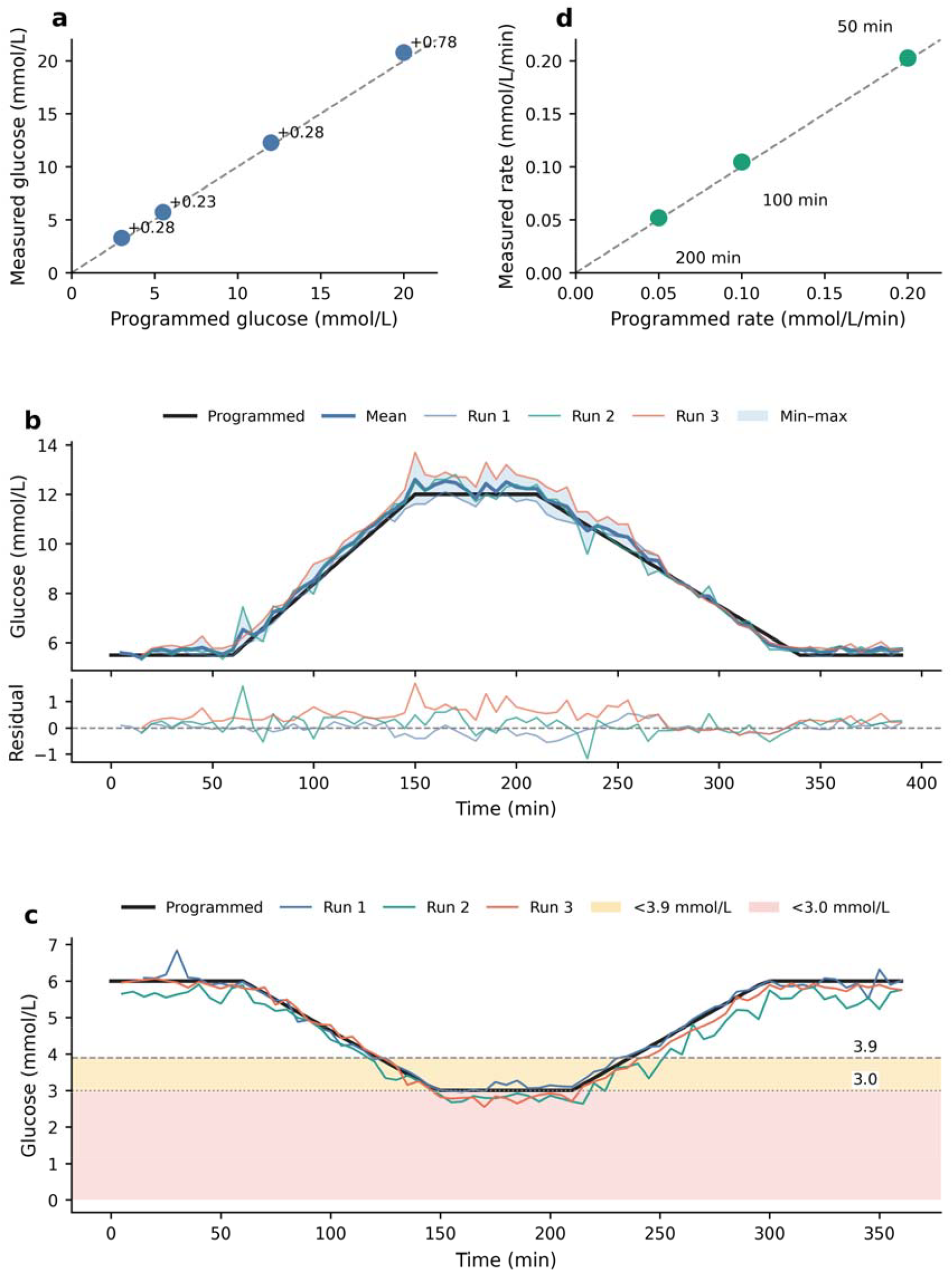
Platform validation: (a) static levels; (b) the 5.5→12.0→5.5 profile showing all runs, the mean curve, range, and residuals; (c) the nocturnal hypoglycemia profile; (d) rate reproduction. A dedicated randomized qualification experiment (four levels, four replicates per control position) confirmed the absence of a statistically significant positional concentration gradient across control positions S1–S6 (one-way ANOVA and Kruskal–Wallis test, all p ≥ 0.20).

The within-series coefficient of variation for these static-level measurements ranged from 0.61% to 2.00%, indicating good short-term repeatability of the reference measurements. To distinguish potential spatial effects from procedural variability, positional homogeneity of the manifold was subsequently evaluated in a dedicated randomized qualification experiment using four programmed concentration levels (3.5, 5.5, 11.0, and 20.0 mmol/L) that did not coincide with the static levels evaluated above. Four replicate measurements were obtained in randomized order at each of the six predefined control positions (S1–S6). Neither one-way ANOVA (F = 0.52–1.34, all p ≥ 0.29) nor the Kruskal–Wallis test (H = 2.41–7.23, all p ≥ 0.20) demonstrated a significant effect of position. Across all concentration levels, the range of position means did not exceed 0.80 mmol/L. These results indicate that the variability observed during the preliminary non-randomized measurements was not confirmed by the subsequent randomized qualification experiment and therefore should not be interpreted as evidence of a stable spatial concentration gradient within the manifold. Under the experimental conditions of the present study, the macrofluidic platform therefore demonstrated practical spatial homogeneity and provided a sufficiently homogeneous reference environment for comparative evaluation of CGM dynamic performance.

### Reproducibility of the 5.5→12.0→5.5 mmol/L Profile

In all three runs the profile preserved the intended sequence of phases: initial plateau, linear rise, upper plateau, linear fall, and return to the initial level. Initial-plateau bias was +0.030, +0.117, and +0.344 mmol/L. The ratio of actual to programmed rise rate was 0.979, 0.978, and 1.083. The first two runs were nearly identical in rise rate, while the third showed moderate acceleration relative to the program.

Amplitude transfer was close to the programmed value but varied between repeated runs. Using the same averaging windows as for the bias calculation (initial plateau 0–60 min and upper plateau 155–210 min), the ratio of realized to programmed amplitude was 0.967 in run 1, 1.022 in run 2, and 1.066 in run 3. The corresponding realized amplitude was 6.287, 6.641, and 6.931 mmol/L against a programmed amplitude of 6.5 mmol/L. Upper-plateau bias was −0.183, +0.258, and +0.775 mmol/L. Pronounced amplitude undershoot was not observed: in the first run the amplitude was 3.3% below the intended value, while in the second and third it was 2.2% and 6.6% above it. This distinction is of fundamental importance for sensor testing: even a moderate deviation of the delivered amplitude from the programmed trajectory could, under direct comparison, be mistakenly attributed to sensor properties.

On the falling phase, the ratio of actual to programmed rate was 0.987, 1.009, and 1.157, and final-plateau bias was +0.077, +0.231, and +0.299 mmol/L. In the first two runs, the falling rate closely matched the program; in the third it was higher. The temporal structure of the residuals was phase-dependent: deviations changed as the trajectory transitioned from plateaus to ramps and could not be removed by a single constant correction. The mean profile and the min–max range shown in Figure 2b demonstrate reproducibility of the overall scenario alongside moderate inter-run differences in amplitude.

### Nocturnal Hypoglycemia Profile

In all three runs, measured GLU crossed the clinically significant threshold of 3.9 mmol/L and reached the range below 3.0 mmol/L. The first crossing of 3.9 mmol/L occurred 117.1–123.3 min after the start of the experiment, suggesting that the time to enter the hypoglycemic range was reproduced fairly consistently. Minimum GLU values were 2.96, 2.64, and 2.55 mmol/L, showing more pronounced variability in the depth of hypoglycemia.

Duration below 3.9 mmol/L was 112.5, 134.6, and 118.8 min. Even more pronounced differences were observed for the range below 3.0 mmol/L: 15.2, 72.6, and 67.5 min. Mean GLU on the low plateau was 3.088, 2.790, and 2.783 mmol/L. Reproducibility of the time to first threshold crossing, in other words, did not guarantee reproducibility of the duration and extent of exposure in the lowest range. For CGM evaluation, this means that each run must be analyzed against its own actually measured trajectory.

After completion of the low plateau, GLU returned to the initial range; recovery was 97.1–98.2% of the initial level. RMSE between the measured and programmed trajectory ranged from 0.162 to 0.388 mmol/L. Overall, the hypoglycemic scenario was reproducible, but the depth and duration of the critical segment remained sensitive to run-specific features (Figure 2c).

### Profiles With Different Rates of Change

At the same programmed amplitude of 5.0→15.0→5.0 mmol/L, transitions of 200, 100, and 50 min were realized. For the 200-min profile, the programmed rate was 0.050 mmol/L/min and the measured rate was 0.0519 mmol/L/min; the measured/programmed ratio was 1.038. Mean upper-plateau GLU was 15.50 mmol/L.

For the 100-min profile, the programmed and measured rates were 0.100 and 0.1045 mmol/L/min, respectively, with a ratio of 1.045 and mean upper-plateau GLU of 15.03 mmol/L. For the 50-min profile, the corresponding values were 0.200 and 0.2027 mmol/L/min, a ratio of 1.013, and mean upper-plateau GLU of 15.35 mmol/L. In all three regimens the actual rate was slightly higher than the programmed rate, but the maximum relative deviation did not exceed 4.5%.

The measured rate of the 100-min profile was 2.01 times that of the 200-min profile, and the rate of the 50-min profile was 1.94 times that of the 100-min profile. The platform therefore reproduced the intended rate ratio of approximately 1:2:4 while maintaining a similar amplitude. This provides a basis for separately investigating the dependence of sensor response on rate of change without simultaneously altering the range of the stimulus (Figure 2d).

### Complex Multiphase Profile

In the complex experiment, the sequence 4→18→3→12→5.5 mmol/L was realized, combining two rises, two falls, and three plateau levels. The measured trajectory preserved the order and direction of all programmed phases but was systematically lower than the program in the middle and upper ranges. The most pronounced discrepancy was observed at the 18 mmol/L plateau: mean measured GLU was 16.20 mmol/L, and the maximum value was 16.8 mmol/L. Consequently, under the conditions of this experiment, the programmed upper level was not reached at any point on the plateau.

Mean phase bias was −0.348 mmol/L on the initial 4 mmol/L plateau, −0.897 mmol/L during the 4→18 rise, −1.800 mmol/L on the 18 mmol/L plateau, −0.790 mmol/L during the 18→3 fall, and −0.284 mmol/L on the 3 mmol/L plateau. In the second part of the profile, bias was −0.883 mmol/L during the 3→12 rise, −1.333 mmol/L on the 12 mmol/L plateau, −0.963 mmol/L during the 12→5.5 fall, and −0.568 mmol/L on the final plateau. The magnitude of the deviation increased at higher concentrations in this experiment, indicating that the difference between programmed and measured GLU was dependent on profile phase and concentration range.

The “measured GLU − programmed” residual curve changed in synchrony with the phases of the profile. On the upper plateaus, the residual became most negative, and its magnitude decreased after return to the low range. This pattern is incompatible with a constant-offset model and shows that a complex profile requires component-wise assessment of amplitude, rate, and phase-dependent deviation (Figure 3).

**Figure 3.**
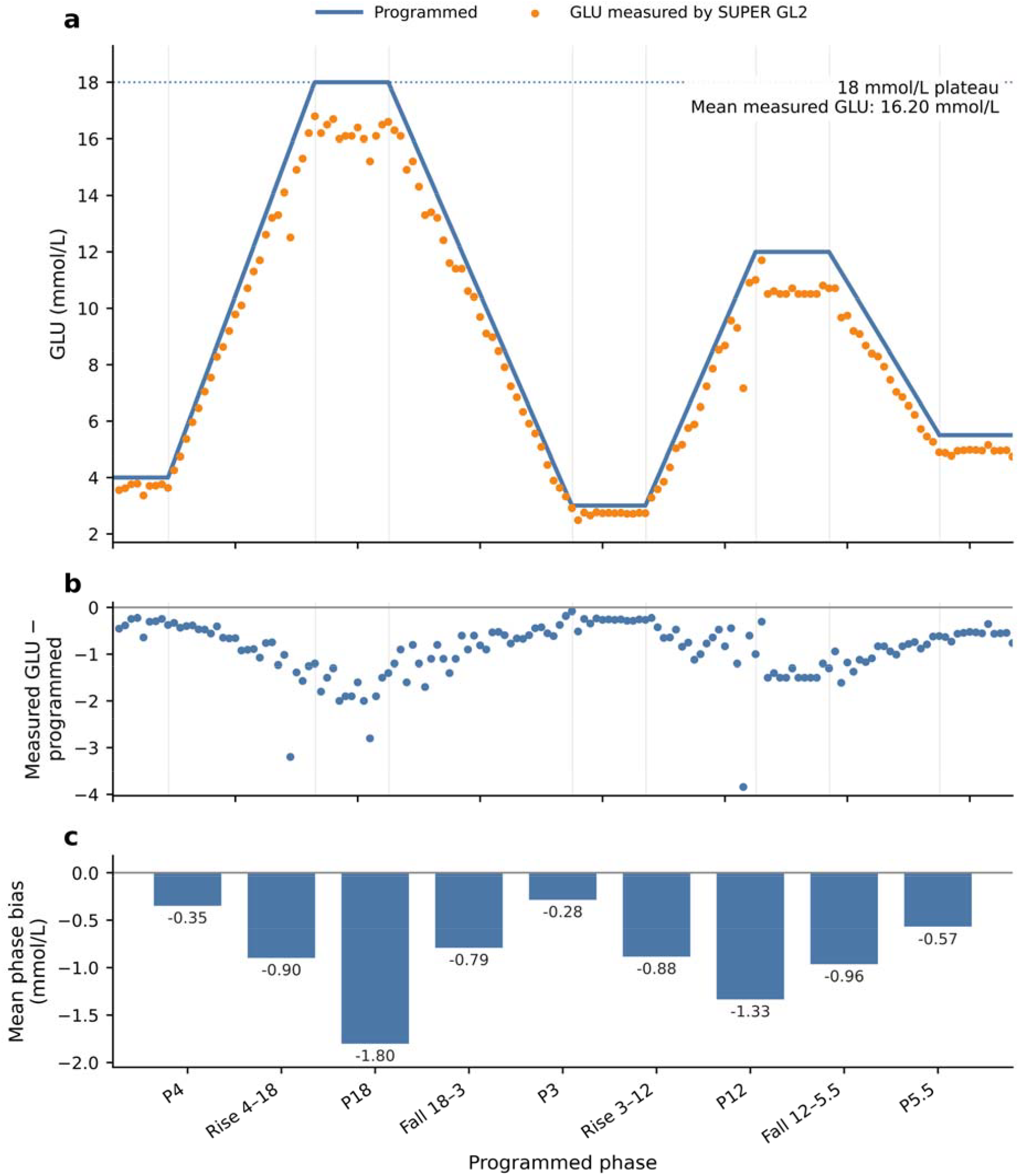
Complex multiphase profile: (a) programmed trajectory and GLU values measured by the SUPER GL2 analyzer; (b) the “measured GLU − programmed” residual over time; (c) mean bias by profile phase.

### Demonstration Characterization of Sensor Dynamic Response

The sensor stage comprised a single sensor of each of two types at 100- and 200-min transitions. Both sensors preserved the direction of GLU change but exhibited opposite types of scale error. Pooled across both regimens, CGM-A had a negative bias of −3.318 mmol/L, MAE of 3.318 mmol/L, RMSE of 3.536 mmol/L, and MARD of 33.70%. The linear-regression slope was 0.656 with a correlation of 0.990. A high correlation together with a slope well below 1 indicated good reproduction of the order of changes alongside pronounced range compression.

For CGM-B, pooled bias was +0.822 mmol/L, MAE was 1.191 mmol/L, RMSE was 1.548 mmol/L, and MARD was 11.49%. Slope was 1.232, and the correlation coefficient was 0.970. CGM-B therefore, on average, overestimated the level and amplified the range of change.

Comparison of the two sensor responses shows that correlation does not reflect the sign of bias or the direction of proportional error.

The amplitude transfer coefficient K_A for CGM-A was 0.682 at the 100-min transition and 0.650 at the 200-min transition, corresponding to consistent amplitude compression of roughly one-third. For CGM-B, K_A was 1.228 and 1.128, showing that amplitude was amplified relative to the actually measured GLU range. Transition duration had a moderate effect on the magnitude of K_A but did not change the direction of scale error for either sensor.

For CGM-A, the rising-rate transfer coefficient K_up was 0.666 and 0.634, and the falling-rate transfer coefficient K_down was 0.730 and 0.626. This means the sensor reproduced only about two-thirds of the actual rate of GLU change. For CGM-B, K_up was 1.564 and 1.276, and K_down was 1.328 and 1.145 — amplification of both rising and falling rate. The difference between K_up and K_down points, further, to a dynamic asymmetry in the response.

After normalization for baseline level and amplitude, normalized RMSE was 0.041 and 0.045 for CGM-A and 0.107 and 0.079 for CGM-B. Normalized correlation was 0.994 and 0.994 for CGM-A and 0.970 and 0.983 for CGM-B. Despite its worse absolute performance, CGM-A therefore reproduced the normalized shape of the profile more accurately. This highlights the distinction between absolute accuracy and dynamic shape fidelity.

Hysteresis loop area was 1.75 and 2.66 (mmol/L)^2^ for CGM-A and 17.81 and 13.33 (mmol/L)^2^ for CGM-B at the 100- and 200-min transitions, respectively. The larger area for CGM-B meant that, at the same GLU value, sensor readings could differ considerably depending on whether concentration was rising or falling. Residual shift after completion of the cycle, calculated using the same averaging windows (initial plateau 0–20 min, final plateau the last 20 min of each run), was −0.37 and +0.13 mmol/L for CGM-A and −0.71 and −0.40 mmol/L for CGM-B at the 100- and 200-min transitions, respectively. For CGM-A, residual shift had opposite signs between the 100- and 200-min regimens, whereas for CGM-B it remained negative in both. No signal loss was recorded; median update interval was 3 min for CGM-A and 5 min for CGM-B.

Time profiles of GLU measured by the SUPER GL2 analyzer and of the sensor readings are shown in Figures 4a and 4b. For visualization, reference SUPER GL2 measurements were connected by linear interpolation only within the observed time range; actual measurement time points are shown as separate markers. CGM-A predominantly retained a negative offset and a compressed range in both regimens. For CGM-B, the sign and magnitude of the error changed between the rise, the upper range, and the fall, consistent with the larger hysteresis loop. Ranking by MARD did not match ranking by normalized RMSE and hysteresis area: the sensor with the smaller absolute error showed the more pronounced dynamic asymmetry.

**Figure 4.**
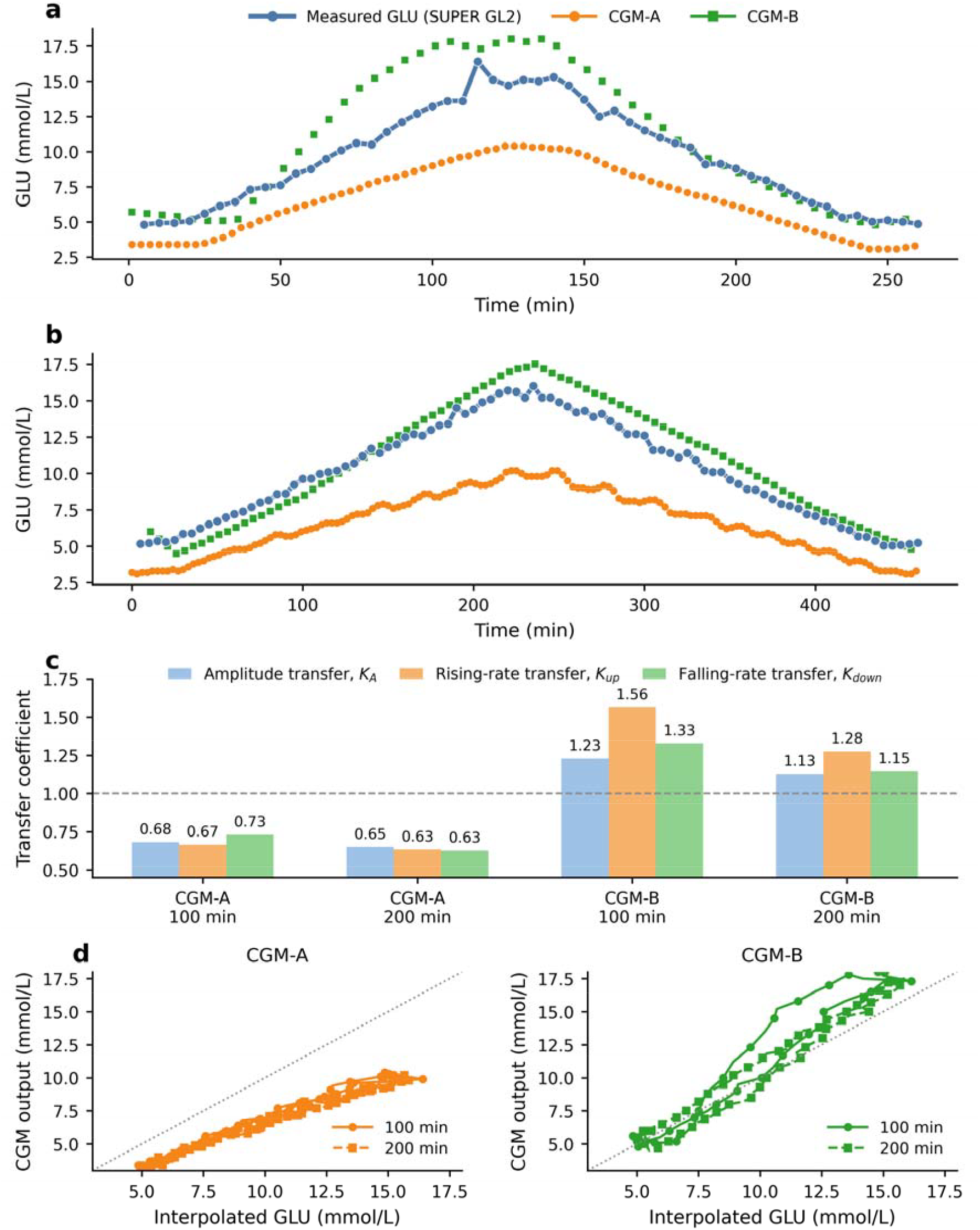
Dynamic characterization of the individual sensors: (a) time profile at the 100-min transition; (b) time profile at the 200-min transition; the blue line shows the linearly interpolated time series of GLU measured by the SUPER GL2 analyzer, and the blue points mark the actual comparator-measurement time points; (c) amplitude transfer coefficient K_A and the rising- and falling-rate transfer coefficients K_up and K_down; (d) hysteresis loops of CGM-A and CGM-B relative to the interpolated GLU series.

## DISCUSSION

The principal finding of this study is that controlled in vitro CGM testing must comprise not one but two sequential measurement tasks. It is first necessary to establish which glucose profile the platform delivered, and only then can the sensor be evaluated against that measured trajectory. In our experiments, the programmed trajectory reproduced the overall scenario and the ratio of rates but did not always achieve the intended amplitude and plateau level. Using the mixing program itself as the direct comparator would therefore have confounded differences between the programmed and realized profile with sensor error.

Good short-term repeatability of the static-level measurements (within-series CV 0.61–2.00%) demonstrates stable performance of the reference measurements. The positional-homogeneity question deserves a further word, since it bears directly on how the static-level bias reported above should be read. Because that qualification experiment used randomized rather than position-blocked sampling, it could distinguish a genuine spatial gradient from ordinary run-to-run scatter — and, as reported in Results, no such gradient emerged at any of the four levels tested. This supports treating the manifold as a single, effectively homogeneous mixing environment: differences observed elsewhere in this study can more confidently be attributed to the sensors under test than to where, geometrically, a given measurement happened to be taken. This conclusion is consistent with the general principle of CGM evaluation whereby the quality of the comparator method, temporal synchronization, and the distribution of paired values substantially affect the resulting accuracy metrics (Kirchsteiger et al., 2015; Pleus et al., 2017; Freckmann et al., 2023). In clinical studies, the comparator method is usually regarded as an external source of uncertainty. In a bench test, the uncertainty of profile generation itself is added to this. The “programmed trajectory — measured glucose — CGM output” analytical cascade is therefore not a graphical device but a necessary structure for result traceability.

The performance of the comparator method itself also warrants separate comment. According to the manufacturer’s operating manual, the coefficient of variation of the SUPER GL2 for repeated measurements at 12 mmol/L is below 1.5%, which is comparable to or better than the typical precision of YSI-class reference glucose analyzers (1–3% CV; Dunseath et al., 2025) and considerably tighter than the ±10% stated in the Russian type-approval certificate (No. 74068-19). The latter figure reflects the regulatory tolerance of a simplified local registration prepared by the distributor for the domestic market, not the instrument’s analytical performance. The coefficient of variation stated by the manufacturer characterizes measurement repeatability rather than systematic agreement with independently prepared reference solutions. Therefore, the static-level differences observed during the initial platform qualification should not be interpreted as evidence of intrinsic platform bias. Instead, they emphasize the importance of independently measuring the realized GLU concentration and using that measured value as the reference for subsequent CGM evaluation. We also note that the manufacturer-specified measurement range (0.5–50 mmol/L) fully spans the entire range covered by this study, including the hypoglycemic profile (minimum 2.55 mmol/L).

The platform reproduced the intended rate ratio of approximately 1:2:4 with a deviation in measured rate of no more than 4.5%. This is a strength of the bench, since it allows the rate of change to be varied at a similar amplitude. In the repeated 5.5→12.0→5.5 profiles, the amplitude transfer coefficient ranged from 0.967 to 1.066, meaning the delivered amplitude was close to, but not identical with, the programmed value. At the same time, the programmed 18 mmol/L plateau was not fully achieved in the complex-profile experiment, illustrating that rate control and level realization should be evaluated as distinct metrological characteristics. Further refinement of the platform may include additional characterization of transient stabilization after switching between programmed concentration levels.

The hypoglycemic profile additionally showed that an identical time to first threshold crossing does not guarantee identical depth and duration of exposure. Duration below 3.0 mmol/L differed more than fourfold between runs. This is of fundamental importance for CGM evaluation, since assessment of recognition time, minimum value, and recovery depends on the actually realized trajectory. Relying on the programmed profile alone could create a false impression of sensor differences where, in fact, the exposure itself differed.

The dependence of CGM accuracy on rate of change is well documented in clinical and laboratory work. During rapid glycemic swings, MARD increases, and the degree of deterioration is system-dependent (Pleus et al., 2015). In vitro studies show that intrinsic lag, filtering, and rate-of-change limiting can produce a substantial discrepancy even in the absence of a physiological delay between blood and interstitial fluid (Davey et al., 2010; Schmelzeisen-Redeker et al., 2015). Our approach complements these observations by quantifying what fraction of the actual amplitude and rate the sensor transmits. This experimental decomposition differs from computational lag-compensation methods such as retrospective deconvolution of the CGM signal (Del Favero et al., 2014): rather than correcting readings after the fact, the proposed bench characterizes the sensor’s intrinsic dynamic error under a controlled input stimulus, which may serve as a basis for subsequently calibrating such algorithms.

The K_A coefficient is interpreted as scaling of the dynamic range. A value below 1 reflects amplitude compression, and a value above 1, amplification. In our proof-of-concept experiments, CGM-A compressed amplitude throughout, whereas CGM-B amplified it. The K_up and K_down coefficients describe rate transfer on the rising and falling phases. Calculating them separately matters because the same full-cycle amplitude can be associated with different behavior on the rise versus the fall. The observed difference between K_up and K_down for CGM-B points to a dynamic asymmetry that a single pooled MARD value would not reveal.

Normalized shape RMSE addresses a different question: after removing offset and scaling, it shows how closely the time-course shape of the sensor profile matches that of measured GLU. CGM-A had markedly worse absolute performance but a lower normalized RMSE. This means its error was, to a greater extent, described by systematic offset and amplitude compression, while the normalized shape was preserved. CGM-B showed better absolute accuracy but more pronounced dynamic distortion. This divergence in ranking confirms the limitations of MARD as a sole criterion (Kirchsteiger et al., 2015; Heinemann et al., 2020; Vigersky and Shin, 2024).

Hysteresis loop area quantitatively characterizes the dependence of a reading on the direction of change. At the same GLU value, a sensor may report different values on the rise versus the fall owing to lag, filtering, membrane dynamics, or algorithmic processing. In our study, the hysteresis loop area for CGM-B exceeded that of CGM-A by a factor of 5 to 10, depending on the regimen, even though CGM-B had the smaller MARD. A low mean relative error therefore does not exclude meaningful phase dependence. Residual shift complemented the hysteresis assessment by characterizing the completeness of the sensor’s return to its initial state after completion of the cycle. For CGM-A its sign differed between the 100- and 200-min regimens, whereas for CGM-B residual shift remained negative in both. These data show that a sensor’s final state after a dynamic stimulus cannot be inferred from MARD, the amplitude transfer coefficient, or hysteresis loop area alone. Residual shift represents an additional characteristic of dynamic sensor behavior that is not reducible to these metrics.

The proposed decomposition is consistent with the general direction of CGM evaluation, moving from a single averaged metric toward a multidimensional performance profile. CG-DIVA assesses the range of expected deviations and accuracy variability (Eichenlaub et al., 2024); error-grid approaches link error to clinical risk (Kovatchev et al., 2004; Klonoff et al., 2024); and reporting recommendations emphasize the need to describe the comparator method, synchronization, and data structure (Freckmann et al., 2023). A controlled dynamic platform adds to these approaches the ability to separately assess amplitude, rate, shape, and hysteresis under predefined scenarios.

Pfützner et al. (2024) demonstrated laboratory dynamic and interference testing of sensors. The present study extends this concept with three elements: mandatory independent measurement of the delivered glucose profile, a set of profiles enabling separate assessment of level and rate, and analytical decomposition of the sensor response. Such an approach could potentially be used during development, comparative technical evaluation, assessment of algorithm changes, and root-cause investigation of nonconformities.

At the same time, the proposed metrics should not yet be regarded as normative acceptance criteria. No generally accepted clinical limits exist for K_A, K_up, K_down, normalized RMSE, or hysteresis area, analogous to those already established for point accuracy under the current CLSI POCT05 guideline (Clinical and Laboratory Standards Institute, 2020). Their reproducibility, uncertainty, and relationship to clinical risk should be investigated across a larger number of sensors and in multiple laboratories. The sensitivity of these metrics to the choice of phase boundaries, interpolation rules, reference-measurement frequency, and smoothing parameters also requires evaluation.

The absence of such limits does not mean the proposed metrics will remain outside a regulatory context. The recent revision of the Diabetes Technology Society Error Grid, supplemented by a separate Trend Accuracy Matrix for assessing errors of rate and direction of change (Klonoff et al., 2024), shows that the field is already moving toward formalized assessment of dynamic, and not only point, accuracy. The metrics proposed in this work decompose dynamic error into components that could give such a matrix concrete technical content: amplitude and rate determine the extent to which trend error is explained by sensor properties rather than by an experimental artifact. For automated insulin delivery systems, where decisions are made in real time based on trend rather than the current value alone (Boughton and Hovorka, 2024), linking a laboratory bench of this kind to clinically oriented risk matrices appears to be the most promising direction for further development of the proposed approach.

Limitations of this study include a single sensor of each type, two rate regimens at the sensor stage, a single complex profile, and incomplete characterization of transient stabilization immediately after switching between programmed flow conditions. The SUPER GL2 manufacturer states a manufacturer-specified measurement range of 0.5–50 mmol/L and a coefficient of variation below 1.5% at 12 mmol/L, which fully covers the range of the present study, including the hypoglycemic profile. The separate Russian instrument registration (type-approval certificate No. 74068-19), prepared by the local distributor with a narrower range (4–30 mmol/L) and a ±10% tolerance, was scoped to the domestic market and should not be read as the manufacturer’s own accuracy specification; in any case, an empirical assessment of reference-measurement uncertainty under the conditions of the present bench was not performed. During the complex multiphase experiment, CGM-A data were recorded in parallel with the SUPER GL2 measurements but were not included in the main analysis, since the original purpose of the complex profile was platform verification rather than sensor characterization; this parallel recording may be regarded as an additional, third proof-of-concept exposure regimen for CGM-A in future analyses. Consecutive time-series points are not independent observations, so a large number of points does not compensate for a small number of sensors. The sensor results should be interpreted strictly as a proof-of-concept of the analytical scheme, not as a comparative evaluation of manufacturers or models.

Despite these limitations, the study shows that bench-based assessment can provide information not available from MARD and correlation alone. The most promising direction is the development of a standardized program that includes profile verification, repeated levels and rates, hypoglycemic scenarios, criteria for temporal synchronization, and predefined rules for calculating dynamic metrics.

## Conclusion

The controlled flow platform enabled realization of static levels, repeated profiles, a hypoglycemic scenario, three rates of change, and a complex trajectory. The programmed profile should be regarded as the experimental setpoint, whereas the independently measured GLU concentration should serve as the reference for evaluation of CGM performance. CGM testing should therefore be structured as the “programmed trajectory — measured glucose — CGM output” cascade. Decomposition into bias, amplitude and rate transfer, shape, timing, and hysteresis provides information that is not available from MARD and correlation alone.

